# Functional properties of CD8+ T cells under lymphopenic conditions

**DOI:** 10.1101/353011

**Authors:** Yu. Yu. Silaeva, A. A. Kalinina, L. M. Khromykh, D. B. Kazansky

**Affiliations:** Federal State Budget Institution of Sciences Institute of Gene Biology, Russian Academy of Sciences, Moscow, Russia; Federal State Budgetary Institution “N.N. Blokhin Medical Research Center of Oncology” of the Ministry of Health of the Russian Federation, Moscow, Russia

## Abstract

A peripheral pool of T lymphocytes consists of several functionally distinct populations of CD8+ T cells. One of the major surface markers that allow to define different populations of T cells are CD44 and CD62L. Expression profile of these markers depends on the functional status of T lymphocyte. Naive CD8+ T cells express CD62L and do not express CD44 (CD62L^hi^CD44^lo^), clones of T cells activated during the primary immune response lose CD62L and express CD44 (CD62L^lo^CD44^hi^). Central memory CD8+ T cells express both CD44 and CD62L (CD44^hi^CD62L^hi^). However expression of activation markers not always correlate with an antigen experience of T cell. It is known, that in lymphopenic conditions peripheral T cells undergo homeostatic proliferation and acquire the memory like surface phenotype CD44^hi^CD62L^hi^, but data about functional activity of these cells remain controversial. We studied relationship between surface expression of the markers CD44 and CD62L and functional properties of CD8+ T cells under lymphopenic conditions. We proved that surface expression of CD44 in not the only condition for T lymphocyte to acquire the functional properties of memory T-cells. It means that identification of CD8 + T-memory cells based solely on an expression profile of surface markers is not completely correct and requires confirmation by functional tests. Moreover, results of our research may become of practical importance for blood transfusion and bone marrow transplantation.

## Introduction

A peripheral pool of T lymphocytes consists of several functionally distinct populations of CD8+ T cells. One of the major surface markers that allow to define different populations of T cells are CD44 and CD62L. Expression of these markers on a surface of T-lymphocytes defines their migration pathways. CD62L mediates interaction of a T-lymphocyte with cells of the high endothelium venules and thus migration of T cells within the lymphoid system. CD44, the receptor for hyaluronic acid, the main component of the extracellular matrix, allows T-lymphocytes to leave the lymphoid system and migrate in the peripheral tissues. Expression profile of these markers varies depending on the functional state of a T-lymphocyte. Naive T cells have the surface phenotype CD62L^hi^CD44^lo^, but CD8 clones activated during the primary immune response lose CD62L expression and acquire surface phenotype CD62L^lo^CD44^hi^. Most CD8 effectors die after antigen clearing and contraction of the immune response, and a small part of them forms a population of long lived memory T cells. The key features of this population are ability to maintain a stable pool for a long time in the absence of a specific antigen and the accelerated immune response to a specific antigen. Long-lived memory CD8 T cells (also called central memory) express both CD44 and CD62L (CD44^hi^CD62L^hi^). However, expression of the surface memory phenotype not always correlates with the real “antigenic experience” of a T cell. It is known, for example, that the pool of peripheral T-lymphocytes of unimmunized gnotobiotic animals, lack of the gut microbiota and kept in sterile conditions, contains virtual memory T-cells specific for the model antigen [1], [2]. It is also known that in lymphopenic conditions, peripheral T-lymphocytes undergo homeostatic proliferation and acquire the surface phenotype of memory T cells CD44^+^CD62L^+^(the so-called “memory-like” CD8+ T cells, T_ML_) [3], [4], [5], [6]. Expression of the memory phenotype by T lymphocytes during homeostatic proliferation is stable and persists for a long time. It was shown that the T_ML_ population cannot downregulate the expression of the activation surface molecules and acquire the naive phenotype [7], [8]. Thus, this population is extremely similar to the true memory T-cells in its phenotypic characteristics. Since the discovery of the T_ML_ population, its functional properties are the main interest for researchers. Despite many studies in this field, information about the functional characteristics of T_ML_ cells remains controversial. Several studies showed that adoptive transfer of naive CD8 + T cells in lymphopenic conditions leads to formation of a T cells population with the memory phenotype, that fully possesses all functional features of true memory cells (accelerated activation and proliferation in response to a specific antigen) [9], [10], but localization and the expression profile of chemokine receptors in this population differs from the expression profile of true memory cells [11]. Some studies revealed a T_ML_ population with immunosuppressive activity [12]. Moreover, it is known that under lymphopenic conditions T cell clones with high affinity to the self MHC molecules (i.e. autoreactive T-cells) proliferate and acquire the memory phenotype [13], [14]. The facts presented above suggest that the surface phenotype of T-lymphocytes may not reflect the actual status of their functioning as naive T-cells and memory cells. This, in turn, can lead to incorrect identification of T-lymphocytes as long-lived memory CD8 + T-cells. Recently our laboratory developed an experimental system that allows to evaluate the immune response of memory T-cells without involving naive T-cells. This experimental system is based on the ability of allospecific memory T cells to proliferate in vitro in response to allogeneic splenocytes or tumor cells subjected to acute heat shock (heating for 1 hour at 45 ° C). It was shown that the primary proliferative response to an alloantigen did not develop if allogeneic antigen-presenting cells were subjected to acute heat shock. The primary allogeneic response block is not canceled by adding exogenous IL-2 in the culture and is not associated with deletion or suppression of T-cell clones capable to respond to an antigen. But long-lived CD8+ memory T cells induced in the primary allogeneic immune response are able to specifically proliferate in response to the allogeneic stimulators exposed to acute heat shock [15], [16]. Since this method allows to register the response of true memory CD8+ T cells previously encountered with a specific antigen, we hypothesized that T cells, whose surface activation phenotype only mimics the phenotype of true memory T cells, would be unable to proliferate in response to stimulators, subjected to acute heat shock. Based on this hypothesis, we investigated the relationship between expression of the surface markers CD44 and CD62L and the functional properties of CD8+ T-lymphocytes populations under lymphopenia conditions. First of all, we investigated allogenic immune response of memory cells in sublethally irradiated immunized and unimmunized mice. Secondly, we studied the allogeneic immune response of splenocytes of sublethally irradiated mice after adoptive transfer of syngeneic splenocytes from intact or immune mice. It was shown that the allogeneic immune response was considerably suppressed in this experimental system despite the fact that adoptively transferred T-cells carried the phenotype of memory like T-cells.

## Materials and methods

### Animals

Mice of the C57BL/6 (K^b^I-A^b^D^b^)), C57BL/10 (K^b^I-A^b^D^b^), B10.D2(R101) (K^d^I-A^d^I-E^d^D^b^), FVB ( K^q^I-A^q^I-E^q^D^q^) and C57BL/6-TgN(ACTbEGFP)1Osb (K^b^I-A^b^D^b^) (hereiafter B6.GFP) strains were obtained from the breeding facility of the Federal State Budgetary Institution “N.N. Blokhin Medical Research Center of Oncology” of the Ministry of Health of the Russian Federation.

### Cell lines

EL4 thymoma cells were i.p. transplanted into syngeneic C57BL/6 mice at 3-5 × 10^6^ per mouse and grown as ascites tumors. In 10–14 days, ascites were collected aseptically by aspiration and washed 3 times by centrifugation in cold phosphate-buffered saline (PBS, pH 7.4) at 4 °C. Viable cells were counted after trypan blue-eosin staining and then used for mice immunization in a dose 2 × 10^7^ cells per mouse.

P815 mastocytoma cells were cultivated *in vitro* in RPMI-1640 medium, supplemented by 10% FBS in concentration between 0,1 – 1×10^6^ cells/ml (37°C, 5% CO_2_). Cells were washed 3 times by centrifugation in cold phosphate-buffered saline (PBS, pH 7.4) at 4 °C. Viable cells were counted after trypan blue-eosin staining and then used for mice immunization in a dose 2 × 10^7^ cells per mouse.

### Immunization

Recombinant B10.D2(R101) mice were immunized intraperitoneally with 1 ml of EL4 cell suspension in PBS (2 × 10^7^ cells per animal). B6.GFP mice were immunized intraperitoneally with 1 ml of P815 cell suspension in PBS (2 × 10^7^ cells per animal). Control non-immunized mice were injected with PBS only. Two months after immunization mice were euthanized by cervical dislocation and their spleens were isolated to prepare cell suspensions.

### Evaluation of tumor growth and rejection

Sublethally irradiated mice of the B10.D2(R101) strain were subcutaneously injected with 0.25 ml of EL-4 lymphoma cell suspension (8 × 10^7^ cells / ml). Then, at 7, 14, and 21 days, the tumor growth was detected by measuring its linear dimensions. The criterion of total tumor rejection was the impossibility of its detection for measuring linear dimensions.

### Mice irradiation

Female B10.D2(R101) and C57BL/6 mice, intact or previously immunized as described above, were irradiated at a dose of 4.5 Gy by the Agat-R therapeutic device (Russia) containing a Co^60^ gamma source with an initial power 1,9×10^14^ Bq. Splenocytes of irradiated mice were used for staining with fluorescent antibodies, and also to determine their proliferative response in a mixed lymphocyte culture (MLR) 10 days after irradiation.

### Cell suspensions

Splenocytes were gently squeezed from each spleen in a Potter homogenizer with a conic pestle in cold PBS at 4 °C and centrifuged (200 g, 5 min). Red blood cells were removed by hypotonic shock and mononuclear cells were washed 3 times by centrifugation in cold PBS at 4 °C. Cells were re-suspended in PBS for subsequent staining with monoclonal antibodies and for adoptive transfer or in the complete medium for a mixed-lymphocyte reaction (MLR). Viable cells were counted after trypan blue-eosin staining.

### Adoptive transfer

Intact B10.D2(R101) or C57BL/6 mice were irradiated at a dose of 4.5 Gy as described above, and 24 hours later they were injected intravenously with 1,5×10^7^ splenocytes from intact or immune animals as described above. 10 days later, splenocytes of the recipient mice were used in MLR as responders.

### Mixed lymphocyte reactions (MLR)

T-lymphocyte proliferation was measured by ^3^H- thymidine incorporation after co-incubation of 4×10^5^ splenocytes with 4×10^5^ mitomycin C-treated (25 mg/ml, 30 min, 37°C) or heat shocked (45°C, 60 min) stimulatory splenocytes in 96-well U-bottom plates. Cells were cultured in a RPMI-1640 medium supplemented with 10% fetal bovine serum (FBS), 4 mM L-glutamine, 20 mM HEPES and 50 mM 2-ME at 37°C, 5% CO_2_ for 72 h. ^3^H- thymidine were added to cells for the last 8 hours of cultivation. The allogeneic response was evaluated by subtraction of the ^3^H- thymidine level in the cell culture with syngeneic stimulatory splenocytes from the radioactive mark level in the culture with allogeneic stimulators.

### Statistical analysis

All data are presented as mean ±SEM. All statistical analyses were performed using a Student’s t test. A p value <0.05 was used to determine statistical significance.

### Staining and analysis of T-lymphocytes on flow cytometer

To analyze T-lymphocyte subpopulations, the following antibodies were used: FITC-conjugated anti-CD8α (Clone 53–6.7), FITC-conjugated anti-CD4 (Clone GK1.5), APC/Cy7-conjugated anti-CD62L (Clone MEL-14), PE-conjugated anti-CD44 (IM7), Alexa Fluor®647- conjugated anti CD3 (Clone 17A2). Cells (1 ×10^6^) were stained with antibodies at 4°C for 35 min. Analysis was performed on the flow cytometer BD FACSCanto II using a FACSDiva 6.0 software (BD Pharmingen). Dead cells were excluded from analysis based on scatter signals and staining with PI ( 7,5·10^−5^ M) or 7AAD Viability Staining Solution (BioLegend). Minimum 1 × 10^6^ events/sample were collected to characterize peripheral T-lymphocyte populations. Further processing of results was performed using Flow Jo 7.6 software (TreeStar Inc., Ashland, OR).

## Results and discussion

The general scheme of the experiments is represented in Fig 1.

**Figure 1.**
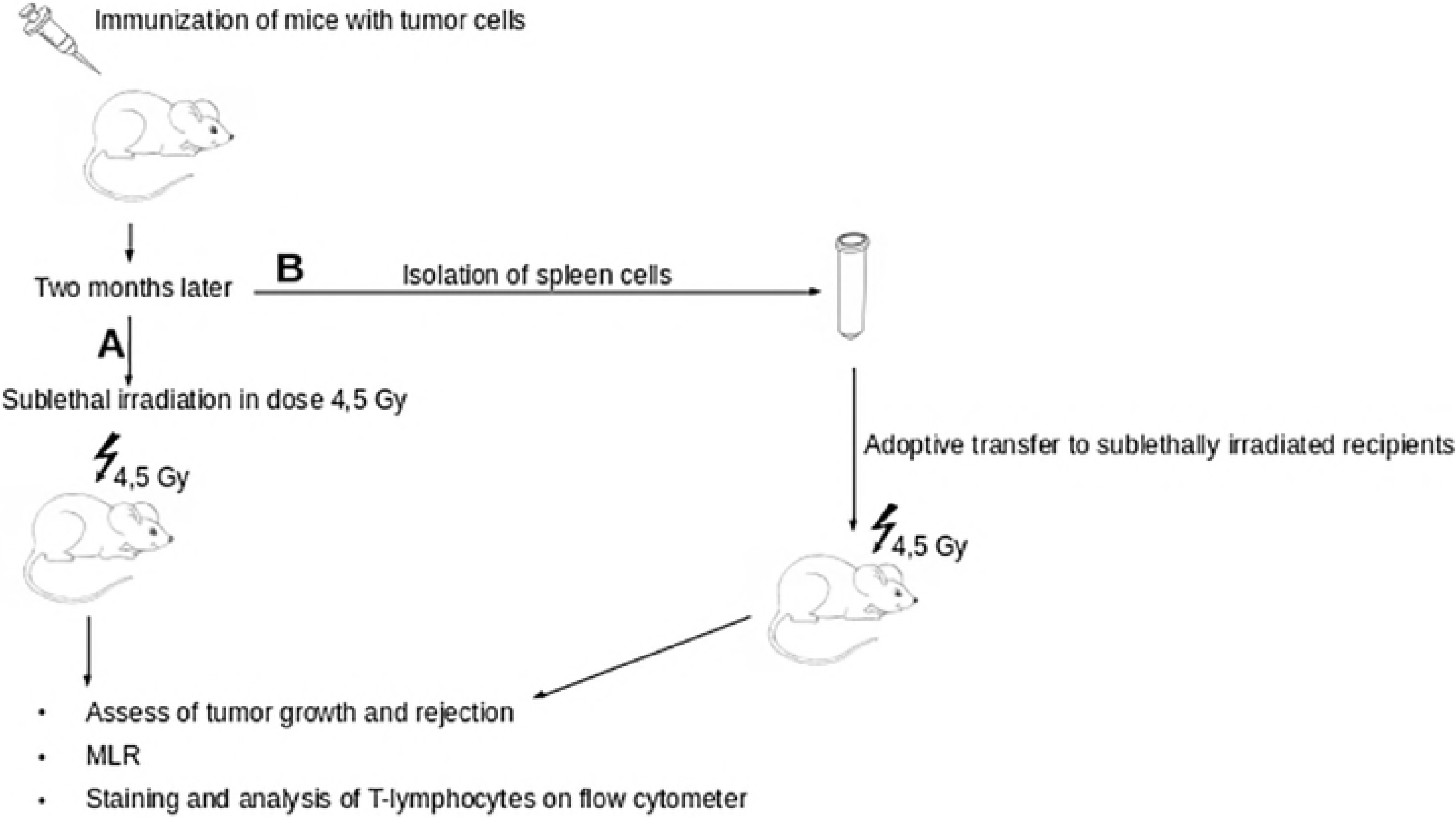
The general scheme of the experiments. (A) - scheme of the experiments without adoptive transfer. (B) - scheme of the experiments adoptive transfer.

### Functional properties and expression profile of surface activation molecules of T-lymphocytes of intact and immunized animals after sublethal irradiation

To study the relationship between expression of CD44 and CD62L and the functional characteristics of peripheral CD8+ T-lymphocytes of intact and immunized mice, the B10.D2(R101) mice were irradiated at a dose of 4.5 Gy, and after 10 days the percentage of naive (CD44-CD62L+), effector (CD44+CD62L−) and central memory cells (CD44+ CD62L +) in the CD8+ population (Figs 1A and 2) as well as the ability of these cells to the primary and secondary allogeneic response in vitro were measured using syngeneic (BL10.D2 (R101) (K^d^I-A^d^I-E^d^D^b^)), allogeneic specific (C57BL /6 (K^b^I-A^b^D^b^)) and allogeneic nonspecific (FVB ( K^q^I-A^q^I-E^q^D^q^)) stimulators (Fig 3). To evaluate the total immune response of naive and memory T cells in vitro, stimulators were preliminarily treated with mitomycin C, and for selective assessment of the memory cell response in vitro stimulators were subjected to acute heat shock (1 hour, 45 ° C). It was shown that after sublethal irradiation the number of CD8+ T cells with the surface phenotype of naive cells is significantly reduced, whereas the proportion of cells with the surface phenotype of effectors and central memory cells increases (Fig 2, lower panel).

**Figure 2.**
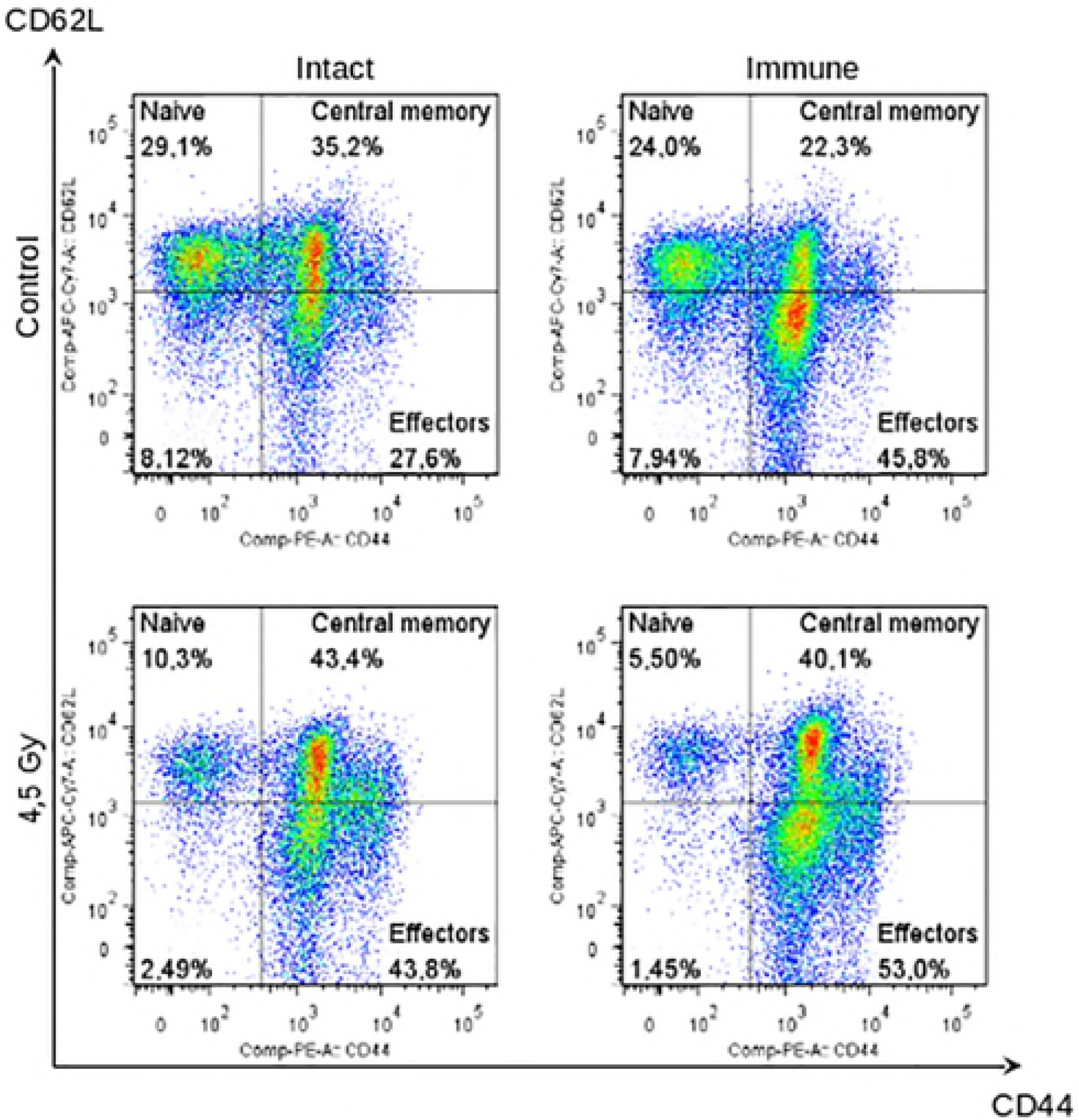
Expression profile of surface activation markers on CD8+ T cells of intact and immune mice after sublethal irradiation. Relative number of CD8+ T cells with phenotype of naive (**Naive**, CD44—CD62L+), effectors (**Effectors**, CD44+CD62L—), and central memory (**Central memory**, CD44+CD62L+) cells in spleens of intact (**left panel, intact**) and immunized (**right panel, immune**) B10.D2(R101) mice 10 days after sublethal irradiation (lower panel, **4,5 Gy).** Data of one representative experiment are given.

The ability of splenocytes to respond to allogeneic stimulators treated with mitomycin C is retained, but somewhat reduced after sublethal irradiation (Fig 3). It is important to note that the ability of splenocytes of pre-immunized irradiated animals to respond to the specific stimulators exposed to acute heat shock increases in comparison with non-irradiated animals. At the same time, the response to nonspecific allogeneic stimulators exposed to acute heat shock remains as low as in unirradiated animals. Thus, in spite of the practically identical ratio of cells with the surface phenotype of naive and activated T-lymphocytes in spleens of intact and immune sublethally irradiated animals (Fig 2), the antigen-specific response of the memory cells was observed only in immunized animals.

**Figure 3.**
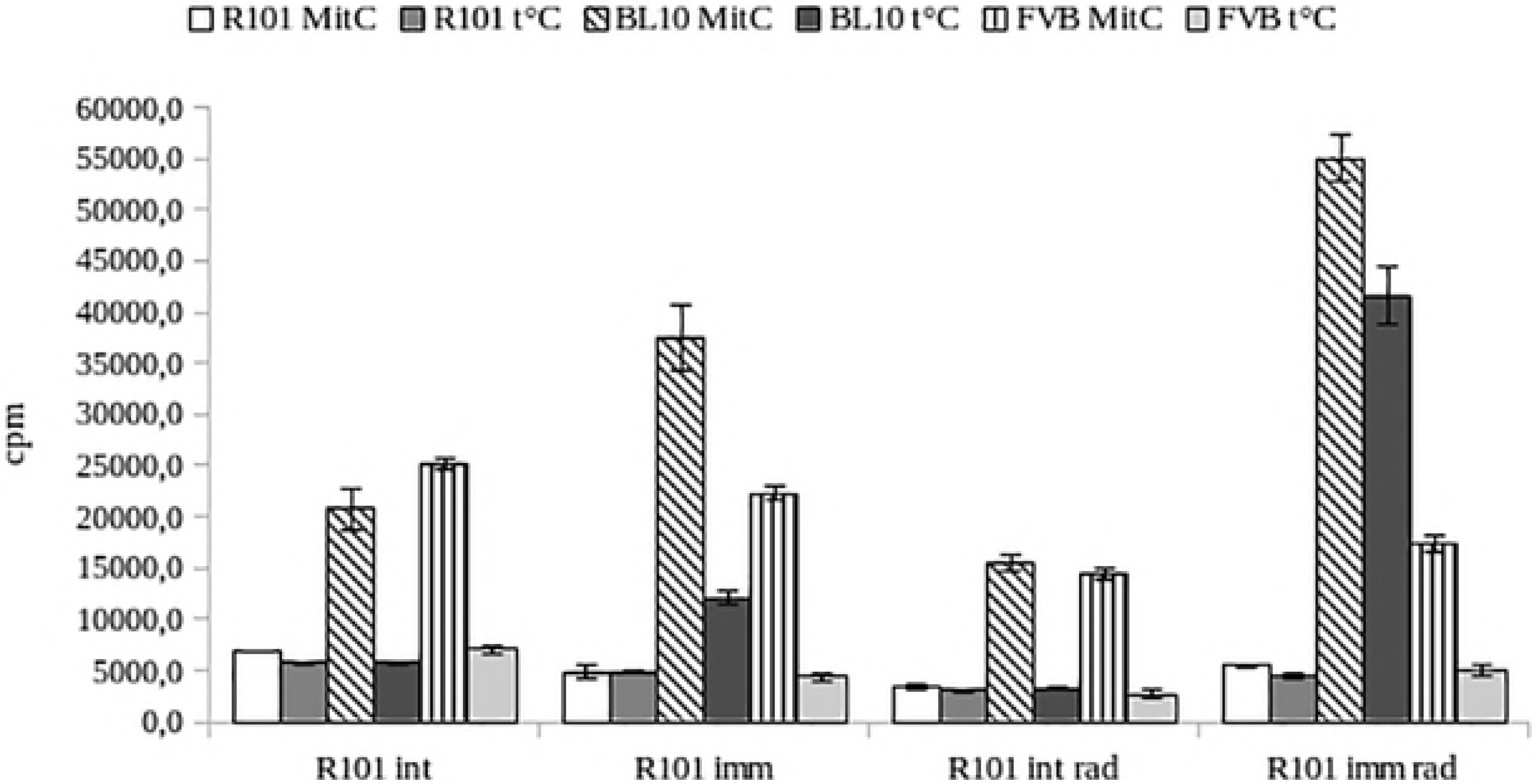
Proliferation in mixed lymphocyte reaction for 72 h of intact and immune B10.D2(R101) mice10 days after sublethal irradiation. Intact (**R101 int**) or immune splenocytes (**R101 imm**) of B10.D2(R101) mice, and intact (**R101 int rad**) or immune splenocytes (**R101 imm rad**) of B10.D2(R101) mice 10 days after sublethal irradiation were used as responders. Splenocytes of B10.D2(R101) (KdDb) mice treated with mitomycin C (**R101 MitC**) or subjected to acute heat shock (1 hour, 45 ° C) (**R101 t°C**); splenocytes of C57BL/10 (KbDb) mice treated with mitomycin C (**BL10 MitC)** or subjected to acute heat shock (**BL10 t°C**); splenocytes of FVB (H-2q) mice treated with mitomycin C (**FVB MitC)** or subjected to acute heat shock (**FVB t°C**) were used as stimulators. Proliferation activity was assessed by incorporation of ^3^H- thymidine for last 8 hours of cultivation. Data of one representative experiment are given.

### Functional properties and surface phenotype of T-lymphocytes of intact and immunized animals after adoptive transfer to sublethally irradiated recipients

To investigate the functional properties of naive cells that acquired the memory phenotype during homeostatic proliferation, sublethally irradiated C57BL/ 6 mice were injected intravenously with 15 × 10^6^ splenocytes of syngeneic intact or immune B6.GFP mice (Fig 1B). Ten days later, 3-5% of GFP+ CD3+ cells from the transgenic GFP expressing donor B6.GFP mice were observed in spleens of the recipients (Fig 4).

**Figure 4.**
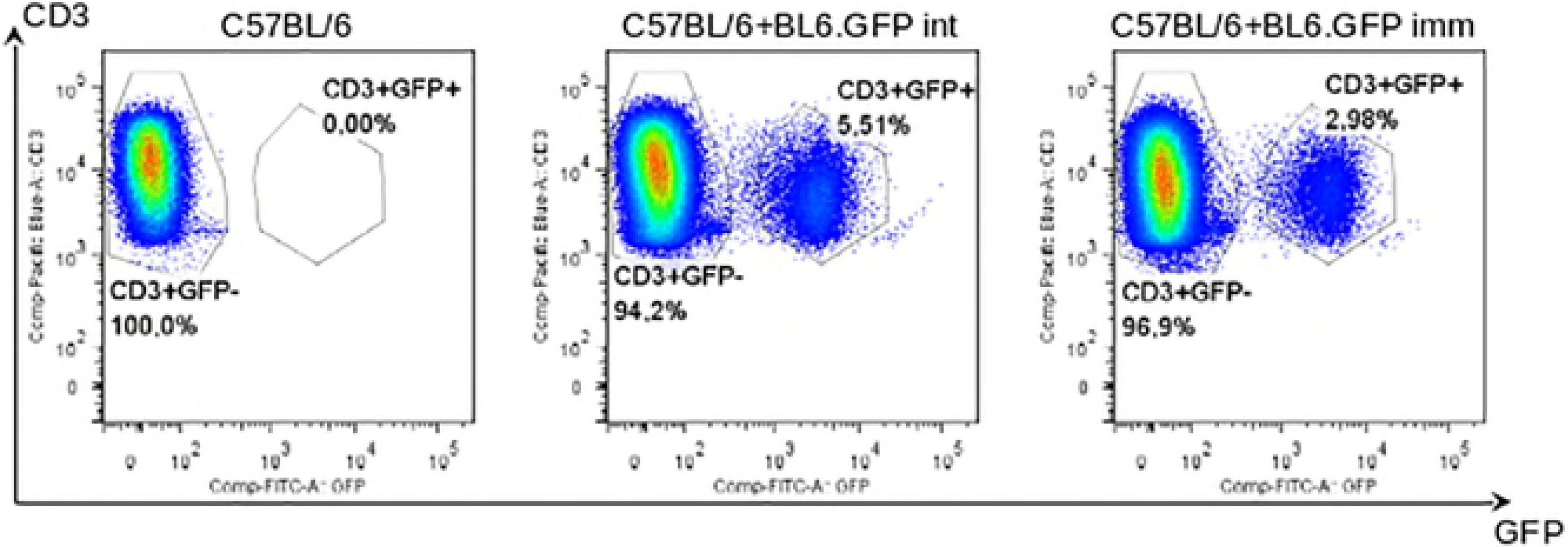
Percentage of GFP^hi^ CD3+ donor cells in spleens of sublethally irradiated recipients. Flow cytometric analysis of GFP^hi^ CD3+ cells in spleens of recipient C57BL/6 mice 10 days after adoptive transfer. Relative number of GFPhiCD3+ **(GFP+)** cells in spleens of intact C57BL/6 mice (**C57BL/6, left panel**), in spleens of sublethally irradiated C57BL/6 mice after adoptive transfer of cells from intact BL6.GFP mice (**C57BL/6+BL6.GFP int, middle panel**), in spleens of sublethally irradiated C57BL/6 mice after adoptive transfer of cells from immune BL6.GFP mice (**C57BL/6+BL6.GFP imm, right panel**). Data for 2,5×10^6^ events. Data of one representative experiment are given.

The ratio of CD4+ and CD8+ cells did not change in comparison with the unirradiated control mice and sublethally irradiated mice without adoptive transfer. The percentage of naive (CD44-CD62L +), effectors (CD44 + CD62L−) and central memory cells (CD44 + CD62L +) in the CD8+ T-cells population of the recipient (GFP−) does not differ from the percentage of these subpopulations in the unirradiated control (Fig 5, left panel, Fig 2, upper panel), but almost all CD8 + donor T cells (GFP +) acquired the activated phenotype (Fig 4, right panel). The ratio of CD44+CD62L− and CD44+CD62L+ populations among CD8+ donor T cells (GFP+) varied within one experiment, but statistically significant difference was not observed in cells number of these subpopulations, adoptively transferred from intact and immunized animals (data not shown).

**Figure 5.**
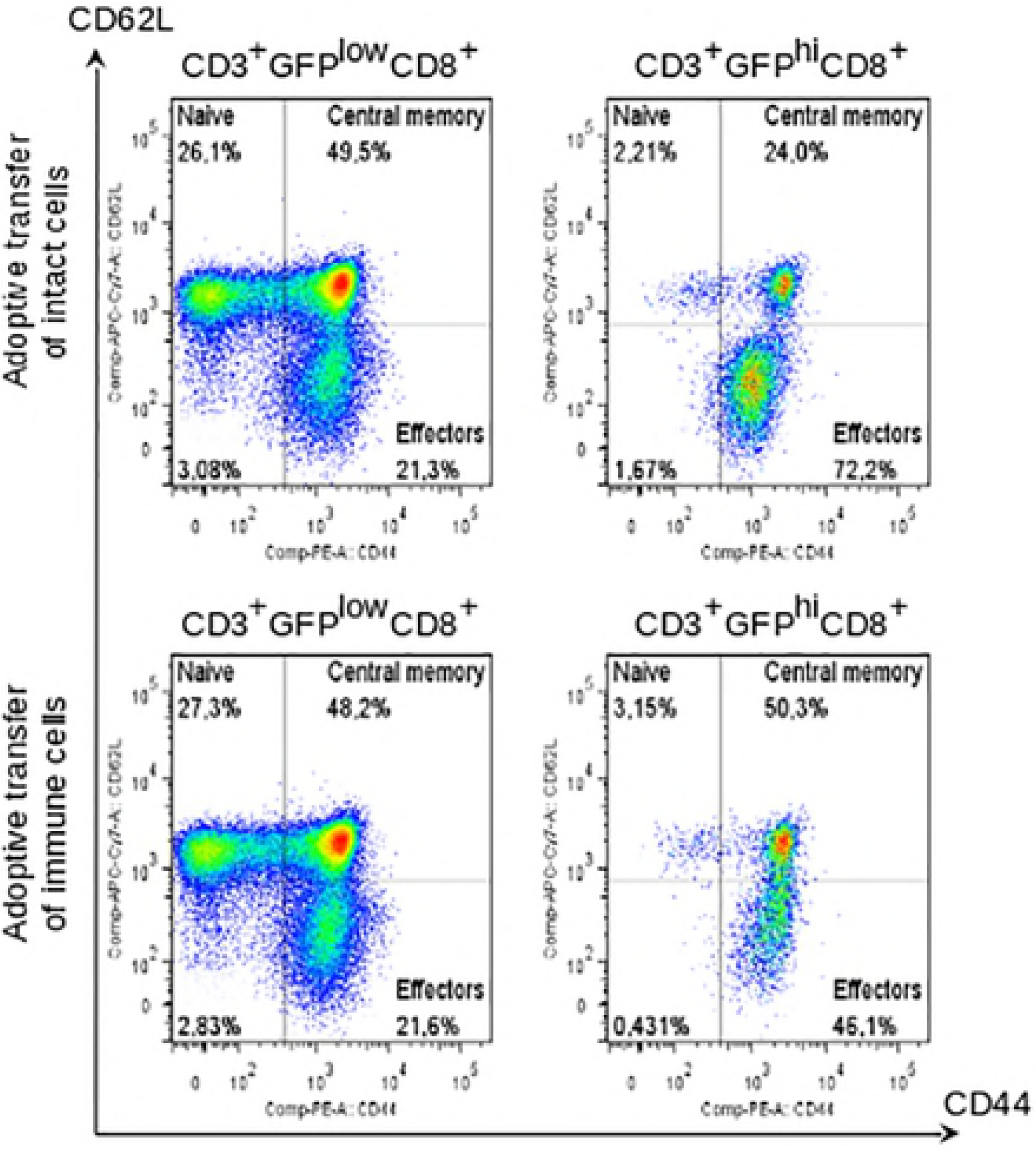
Expression profile of surface activation markers on CD8+ T cells of donor mice and recipients after adoptive transfer. Relative number of CD8+ T cells with phenotype of naive (**Naive**, CD44—CD62L+), effectors (**Effectors**, CD44+CD62L—), and central memory (**Central memory**, CD44+CD62L+) cells in spleens of sublethally irradiated recipients 10 days after adoptive transfer. **CD3+GFPlowCD8**+ - flow cytometric analysis of recipient cells; **CD3+GFPhiCD8**+ - flow cytometric analysis of donor cells; **Upper panel** - adoptive transfer of intact cells; **Lower panel**- adoptive transfer of immune cells. Data of one representative experiment are given.

We investigated a proliferative response in vitro of splenocytes of sublethally irradiated mice after adoptive transfer of spleen cells from syngeneic intact or immune animals (Fig. 6). It was shown that the primary and secondary immune responses in vitro of the recipients after adoptive transfer of intact or immune splenocytes are practically absent. Next, we studied the ability of sublethally irradiated B10.D2 (R101) mice to reject allogeneic EL-4 lymphoma cells after adoptive transfer. Animals were subcutaneously injected with 20 × 10^6^ EL-4 lymphoma cells simultaneously with the adoptive transfer of syngeneic splenocytes, and tumor growth and rejection were subsequently observed (Table 1). It was shown that the rejection of EL-4 lymphoma in irradiated mice after adoptive transfer is much slower than in irradiated animals without adoptive transfer, i.e. adoptive transfer of syngeneic splenocytes from immune and intact animals does not accelerate, but even slows down the process of tumor rejection. Thus, although splenocytes of both intact and immunized mice after adoptive transfer to sublethally irradiated animals acquired the surface phenotype of activated T cells their primary and secondary immune responses in vitro and in vivo are significantly suppressed.

**Figure 6.**
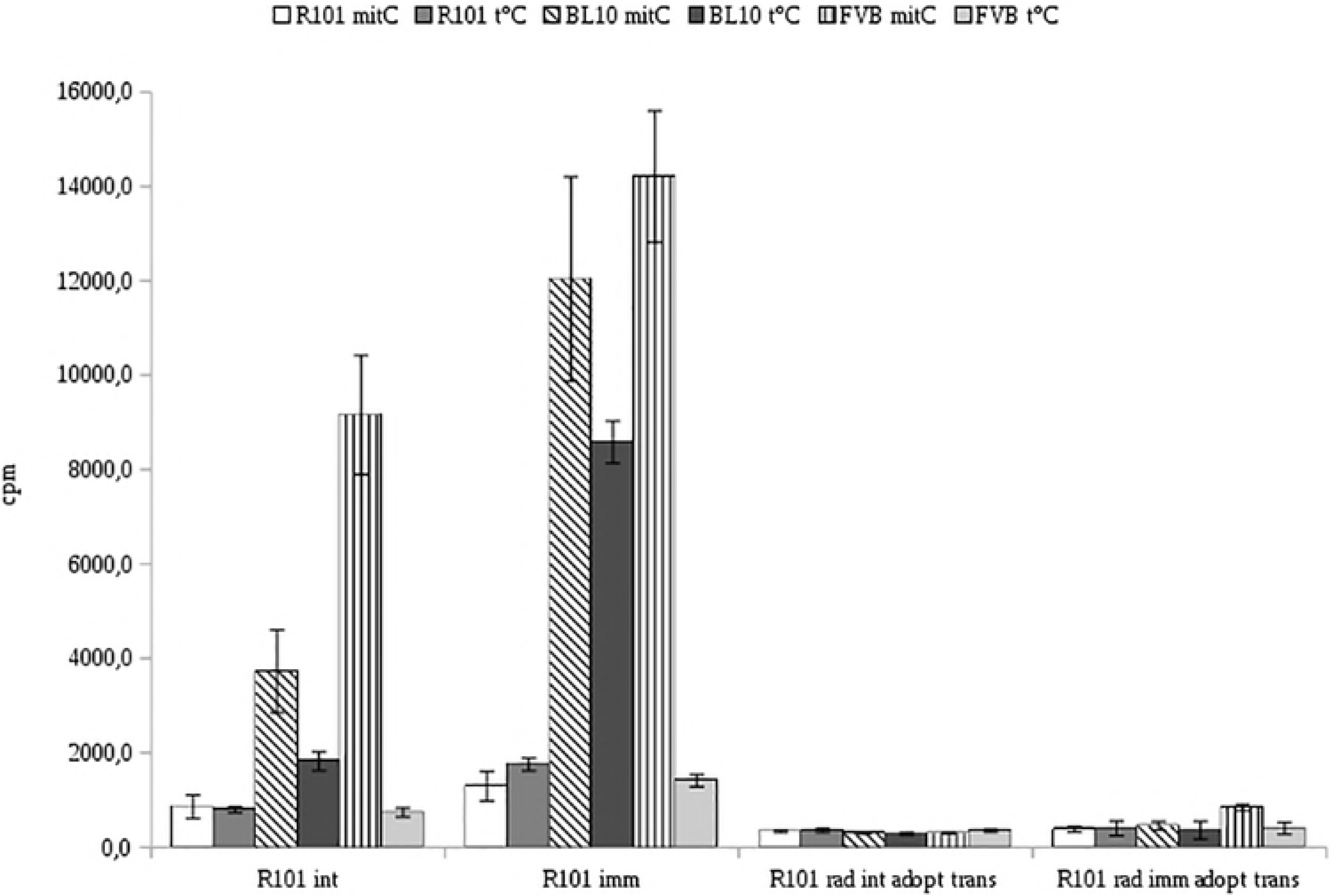
Allogeneic immune response of splenocytes of sublethally irradiated mice after adoptive transfer. Proliferation in mixed lymphocyte reaction for 72 h of splenocytes of sublethally irradiated B10.D2(R101) mice 10 days after adoptive transfer of intact or immune syngeneic splenocytes. Intact (**R101 int**) or immune splenocytes (**R101 imm**) of B10.D2(R101) mice, splenocytes of sublethally irradiated mice after adoptive transfer of intact (**R101 rad int adopt trans**) or immune (**R101 rad int adopt trans**) cells of B10.D2(R101) mice were used as responders. Splenocytes of B10.D2(R101) (KdDb) mice treated with mitomycin C (**R101 MitC)** or subjected to acute heat shock (1 hour, 45 ° C) (**R101 t°C**); splenocytes of C57BL/10 (KbDb) mice treated with mitomycin C (**BL10 MitC)** or subjected to acute heat shock (**BL10 t°C**); splenocytes of FVB (H-2q) mice treated with mitomycin C (**FVB MitC)** or subjected to acute heat shock (**FVB t°C**) were used as stimulators. Proliferation activity was assessed by incorporation of ^3^H- thymidine for last 8 hours of cultivation. Data of one representative experiment are given.

**Table 1.** Evaluation of tumor growth and rejection after subcutaneous injection of 2 × 10^7^ EL-4 lymphoma cells

|  | Tumor growth, days post subcutaneous injection |  |  |
| --- | --- | --- | --- |
|  | 7 days | 14 days | 21 days |
| B10.D2(R101) intact (n=3) | + | — | — |
| B10.D2(R101) immune (n=3) | — | — | — |
| B10.D2(R101) intact sublethally irradiated (n=3) | + | — | — |
| B10.D2(R101) immune sublethally irradiated (n=3) | — | — | — |
| B10.D2(R101) sublethally irradiated after adoptive transfer of intact cells (n=3) | + | + | — |
| B10.D2(R101) sublethally irradiated after adoptive transfer of immune cells (n=3) | + | + | — |

Until recently, it was considered that the phenotype of memory cells one-to-one corresponds to their functional properties, i.e. the ability to prolong self-sustainability, resistance to apoptosis, simplified activation conditions, accelerated proliferation and acquisition of effector functions in response to a specific antigen. However, the recent data indicate that there is no rigid relationships between a surface activation phenotype and functional characteristics of a cell. So, the memory-like population formed under the lymphopenic conditions is not always possesses the functions of memory cells, moreover, it was shown that in some cases the population of CD8+CD122+CD44+ cells has immunosuppressive activity [17], [12]. Thus it is arguable if identification of CD8+ T-memory cells based solely on the surface phenotype is absolutely correct. As expected, we showed that CD8 + T-lymphocytes of immune and intact sublethally irradiated mice acquired the surface phenotype CD44+. However, T cells of nonimmunized sublethally irradiated animals did not proliferate in response to allogeneic stimulators exposed to acute heat shock. After adoptive transfer of splenocytes to sublethally irradiated animals, donor T lymphocytes also acquire the memory-like phenotype. But an allogeneic immune response of these cells in vitro is significantly suppressed. Moreover, the specific response of memory T-cells to the allogeneic stimulators subjected to acute heat shock is not observed even after the adoptive transfer of splenocytes from immunized animals. In our experimental systems, lymphopenia was caused by sublethal irradiation. It is known that memory cells are less prone to apoptosis then naive T cells [18], [19], express more bcl-2 [20], [21] and, as a result, are more radioresistant. Thus, after sublethal irradiation spleen cells of immunized animals can be enriched with the true memory cells due to both homeostatic proliferation and death of naive cells. This can explain stimulation of the secondary immune response in sublethally irradiated immune animals. After adoptive transfer of syngeneic splenocytes to sublethally irradiated animals, the proliferative immune response (both the primary and the secondary) is almost completely absent. he lack of response of adoptively transferred immune splenocytes to stimulators exposed to acute heat shock may be explained by dilution of the true memory T cells pool during homeostatic proliferation [22]. This phenomenon may be due to the fact that naive T cells proliferate much faster than memory cells under lymphopenic conditions. Although during such proliferation naive T cells acquire the surface phenotype of effectors and memory cells, they only phenotypically imitate the true memory cells. Much more mysterious is the lack of the primary immune response in this experimental system. It is known that T-cell clones with high affinity to the self MHC/peptide complexes have the highest proliferative activity under the lymphopenic conditions [23], [24]. Thus, after adoptive transfer of splenocytes in the lymphopenic conditions, autoreactive T cell clones not capable to the allogeneic immune response, can actively proliferate [25]. Other possible explanations for the observed phenomenon may be the formation of a suppressor population from the donor T-cells after adoptive transfer to an irradiated animal [12] or the cytotoxic activity of the TML population.

## Conclusions

Thus, it is clear that expression of CD44 on the surface of a T cell does not lead to automatic acquisition of the functional properties of memory T cells. This means that identification of CD8+ T-memory cells based solely on the surface phenotype is not completely correct and requires confirmation by functional tests. Moreover, results of our research may become of practical importance for blood transfusion and bone marrow transplantation, as the memory-like CD8+ T cells population is likely to exist in a human organism [26].

## References

1. Haluszczak C, Akue AD, Hamilton SE, Johnson LDS, Pujanauski L, Teodorovic L, et al. The antigen-specific CD8 ^+^ T cell repertoire in unimmunized mice includes memory phenotype cells bearing markers of homeostatic expansion. J Exp Med. 2009;206: 435–448. doi:10.1084/jem.20081829

2. White JT, Cross EW, Kedl RM. Antigen-inexperienced memory CD8+ T cells: where they come from and why we need them. Nat Rev Immunol. 2017;17: 391–400. doi:10.1038/nri.2017.34

3. Cho BK, Rao VP, Ge Q, Eisen HN, Chen J. Homeostasis-stimulated proliferation drives naive T cells to differentiate directly into memory T cells. J Exp Med. 2000;192: 549–556.

4. Goldrath AW, Bogatzki LY, Bevan MJ. Naive T cells transiently acquire a memory-like phenotype during homeostasis-driven proliferation. J Exp Med. 2000;192: 557–564.

5. Murali-Krishna K, Ahmed R. Cutting edge: naive T cells masquerading as memory cells. J Immunol Baltim Md 1950. 2000;165: 1733–1737.

6. Jameson SC, Lee YJ, Hogquist KA. Innate Memory T cells. Advances in Immunology. Elsevier; 2015. pp. 173–213. doi:10.1016/bs.ai.2014.12.001

7. Ge Q, Hu H, Eisen HN, Chen J. Naïve to memory T-cell differentiation during homeostasis-driven proliferation. Microbes Infect. 2002;4: 555–558.

8. Tanchot C, Le Campion A, Martin B, Léaument S, Dautigny N, Lucas B. Conversion of naive T cells to a memory-like phenotype in lymphopenic hosts is not related to a homeostatic mechanism that fills the peripheral naive T cell pool. J Immunol Baltim Md 1950. 2002;168: 5042–5046.

9. Moxham VF, Karegli J, Phillips RE, Brown KL, Tapmeier TT, Hangartner R, et al. Homeostatic proliferation of lymphocytes results in augmented memory-like function and accelerated allograft rejection. J Immunol Baltim Md 1950. 2008;180: 3910–3918.

10. Oghumu S, Terrazas CA, Varikuti S, Kimble J, Vadia S, Yu L, et al. CXCR3 expression defines a novel subset of innate CD8 ^+^ T cells that enhance immunity against bacterial infection and cancer upon stimulation with IL-15. FASEB J. 2015;29: 1019–1028. doi:10.1096/fj.14-264507

11. Cheung KP, Yang E, Goldrath AW. Memory-Like CD8+ T Cells Generated during Homeostatic Proliferation Defer to Antigen-Experienced Memory Cells. J Immunol. 2009;183: 3364–3372. doi:10.4049/jimmunol.0900641

12. Wang L-X, Li Y, Yang G, Pang P, Haley D, Walker EB, et al. CD122+CD8+ Treg suppress vaccine-induced antitumor immune responses in lymphodepleted mice. Eur J Immunol. 2010;40: 1375–1385. doi:10.1002/eji.200839210

13. Le Campion A, Gagnerault M-C, Auffray C, Becourt C, Poitrasson-Riviere M, Lallemand E, et al. Lymphopenia-induced spontaneous T-cell proliferation as a cofactor for autoimmune disease development. Blood. 2009;114: 1784–1793. doi:10.1182/blood-2008-12-192120

14. White JT, Cross EW, Burchill MA, Danhorn T, McCarter MD, Rosen HR, et al. Virtual memory T cells develop and mediate bystander protective immunity in an IL-15-dependent manner. Nat Commun. 2016;7: 11291. doi:10.1038/ncomms11291

15. Kazanskiĭ DB, Petrishchev VN, Shtil’ AA, Chernysheva AD, Sernova NV, Abronina IF, et al. [Use of heat shock of antigen-presenting cells for functional testing of allospecificity memory T-cells]. Bioorg Khim. 1999;25: 117–128.

16. Pobezinskaya EL, Pobezinskii LA, Silaeva YY, Anfalova TV, Khromykh LM, Tereshchenko TS, et al. Cross reactivity of T cell receptor on memory CD8+ cells isolated after immunization with allogeneic tumor cells. Bull Exp Biol Med. 2004;137: 493–498.

17. Suzuki H, Shi Z, Okuno Y, Isobe K. Are CD8+CD122+ cells regulatory T cells or memory T cells? Hum Immunol. 2008;69: 751–754. doi:10.1016/j.humimm.2008.08.285

18. Grayson JM, Harrington LE, Lanier JG, Wherry EJ, Ahmed R. Differential sensitivity of naive and memory CD8+ T cells to apoptosis in vivo. J Immunol Baltim Md 1950. 2002;169: 3760–3770.

19. Yao Z, Jones J, Kohrt H, Strober S. Selective Resistance of CD44 ^hi^ T Cells to p53-Dependent Cell Death Results in Persistence of Immunologic Memory after Total Body Irradiation. J Immunol. 2011;187: 4100–4108. doi:10.4049/jimmunol.1101141

20. Grayson JM, Zajac AJ, Altman JD, Ahmed R. Cutting edge: increased expression of Bcl-2 in antigen-specific memory CD8+ T cells. J Immunol Baltim Md 1950. 2000;164: 3950–3954.

21. Kurtulus S, Tripathi P, Moreno-Fernandez ME, Sholl A, Katz JD, Grimes HL, et al. Bcl-2 Allows Effector and Memory CD8+ T Cells To Tolerate Higher Expression of Bim. J Immunol. 2011;186: 5729–5737. doi:10.4049/jimmunol.1100102

22. Peacock CD, Kim S-K, Welsh RM. Attrition of virus-specific memory CD8+ T cells during reconstitution of lymphopenic environments. J Immunol Baltim Md 1950. 2003;171: 655–663.

23. Ge Q, Rao VP, Cho BK, Eisen HN, Chen J. Dependence of lymphopenia-induced T cell proliferation on the abundance of peptide/ MHC epitopes and strength of their interaction with T cell receptors. Proc Natl Acad Sci. 2001;98: 1728–1733. doi:10.1073/pnas.98.4.1728

24. Hao Y, Legrand N, Freitas AA. The clone size of peripheral CD8 T cells is regulated by TCR promiscuity. J Exp Med. 2006;203: 1643–1649. doi:10.1084/jem.20052174

25. Villa A, Marrella V, Rucci F, Notarangelo LD. Genetically determined lymphopenia and autoimmune manifestations. Curr Opin Immunol. 2008;20: 318–324. doi:10.1016/j.coi.2008.02.001

26. Van Kaer L. Innate and virtual memory T cells in man. Eur J Immunol. 2015;45: 1916–1920. doi:10.1002/eji.201545761

